# Maternal antioxidant treatment prevents behavioural and neural changes in offspring exposed to prenatal social stress

**DOI:** 10.1101/701292

**Authors:** H Scott, TJ Phillips, Y Sze, A Alfieri, MF Rogers, CP Case, PJ Brunton

**Affiliations:** School of Clinical Sciences, University of Bristol, Learning & Research Building, Southmead Hospital, Bristol BS10 5NB, UK; UK Dementia Research Institute, Cardiff University, Hadyn Ellis Building, Maindy Road, Cardiff CF24 4HQ, UK; Division of Neurobiology, The Roslin Institute, University of Edinburgh, Easter Bush, Midlothian EH25 9RG, UK; Centre for Discovery Brain Sciences, University of Edinburgh, Hugh Robson Building, 15 George Square, Edinburgh EH8 9XD, UK; Intelligent Systems Laboratory, University of Bristol, Merchant Venturers Building, Woodland Road, Bristol BS8 1UB, UK; Zhejiang University-University of Edinburgh Joint Institute, Zhejiang University School of Medicine, International Campus, Haining, Zhejiang 314400, P.R. China.

## Abstract

Maternal exposure to social stress during pregnancy is associated with an increased risk of psychiatric disorders in the offspring in later life. However, the mechanism through which the effects of maternal stress are transmitted to the foetus is unclear. Using a rat model, we explored the mechanisms by which maternal social stress is conveyed to the foetus and the potential for targeted treatment to prevent disease in the offspring. Maternal stress increased circulating corticosterone in the mother, but not the foetuses. Maternal stress also induced oxidative stress in the placenta, but not in the foetal brain, and this was prevented by administration of a nanoparticle-bound antioxidant. Moreover, antioxidant treatment prevented prenatal stress-induced anxiety-like behaviour in the adult male offspring, along with several stress-induced neuroanatomical, neurochemical and gene expression changes in the offspring brain. Importantly, many of these neural effects were mimicked in neuronal cultures by application of placental-conditioned medium or foetal plasma from stressed pregnancies. Both placental-conditioned medium and foetal plasma contained differentially abundant extracellular microRNAs following prenatal stress. The present study highlights the crucial role of the placenta, and the molecules it secretes, in foetal brain development and provides evidence of the potential for treatment that can prevent maternal stress-induced foetal programming of neurological disease.

## Introduction

Stress and anxiety experienced during pregnancy are associated with an increased risk of psychiatric disorders, such as anxiety and depression, attention deficit-hyperactivity disorder, autism spectrum disorder and schizophrenia^1–3^, in the offspring in later life. In a rodent model, maternal social stress in pregnancy gives rise to physiological and behavioural changes in the offspring, including low birthweight, abnormal glucose homeostasis, heightened anxiety behaviour, enhanced stress responses, cognitive deficits and abnormal social behaviours^4–6^. Despite much research on the consequences of maternal stress on the offspring, it is still unclear how the effects of psychological stress are transmitted from mother to foetus. Most rodent studies have focussed on a role for maternal glucocorticoids as a mediating factor, with mixed results^7,8,9^. Moreover, in women maternal stress or anxiety during pregnancy is only weakly associated with raised maternal cortisol, if at all^7^.

The placenta, as the interface between mother and foetus, is likely to play a key role in maternal programming of foetal development^10–12^. Indeed, maternal exposure to different stressors affects placental growth and morphology^13,14^, metabolism and endocrine function^15–18^, gene expression^15,16, 19–23^ and the epigenetic profile^23–25^ – characteristics that have been associated with altered foetal neurodevelopment and disease^22,26–30^. Moreover, our recent research suggests that unknown ‘factors’ released from the placenta can induce neurological changes in the offspring of rats exposed to hypoxia during pregnancy^31^. Maternal treatment that reduces oxidative stress in the placenta, but does not directly affect the foetus, is able to prevent abnormal neurodevelopmental changes in the offspring induced by maternal hypoxia^31^.

It is not known if the placenta plays a role in mediating the effects of maternal social stress on foetal neurodevelopment and adult behavioural phenotypes. Moreover, it is not known if *in utero* intervention through treatment of placental changes can rescue altered neurodevelopment in the offspring and hence prevent abnormal behaviour in later life. Here, we used a rat model to examine the role of the placenta in mediating the effects of maternal social stress on the foetus. After ascertaining a role for maternal social stress in inducing oxidative stress in the placenta, we tested if maternal treatment with a nanoparticle-bound antioxidant is able to prevent both the short- and long-term neurological and behavioural effects of prenatal social stress in the offspring. Given dysfunction of the GABAergic, glutamatergic and dopaminergic systems have been implicated in the pathophysiology of psychiatric disorders for which prenatal stress increases risk^32–35^, we hypothesised that maternal exposure to social stress during pregnancy may lead to imbalances in the expression of GABA and glutamate receptors, along with neuroanatomical changes, in particular of tyrosine hydroxylase-immunoreactive dopaminergic neurons and parvalbumin-positive GABAergic interneurons.

## Results

### 1. Maternal social stress increases circulating corticosterone in the mother but not in the foetus

Stress-induced elevations in maternal glucocorticoids have been proposed to mediate the effects of prenatal stress on the offspring; however, there is limited evidence to support this assertion^8^. Circulating corticosterone concentrations were significantly greater in mothers exposed to repeated social stress during late pregnancy, compared with non-stressed mothers, irrespective of drug treatment (Figure 1a). In contrast, neither maternal social stress nor drug treatment had any significant effect on plasma corticosterone concentrations in the male (Figure 1b) or female foetuses (Figure 1c). These data suggest that the programming effects of prenatal social stress are unlikely to be mediated directly by changes in foetal corticosterone levels and indicate other factors may be involved.

**Figure 1.**
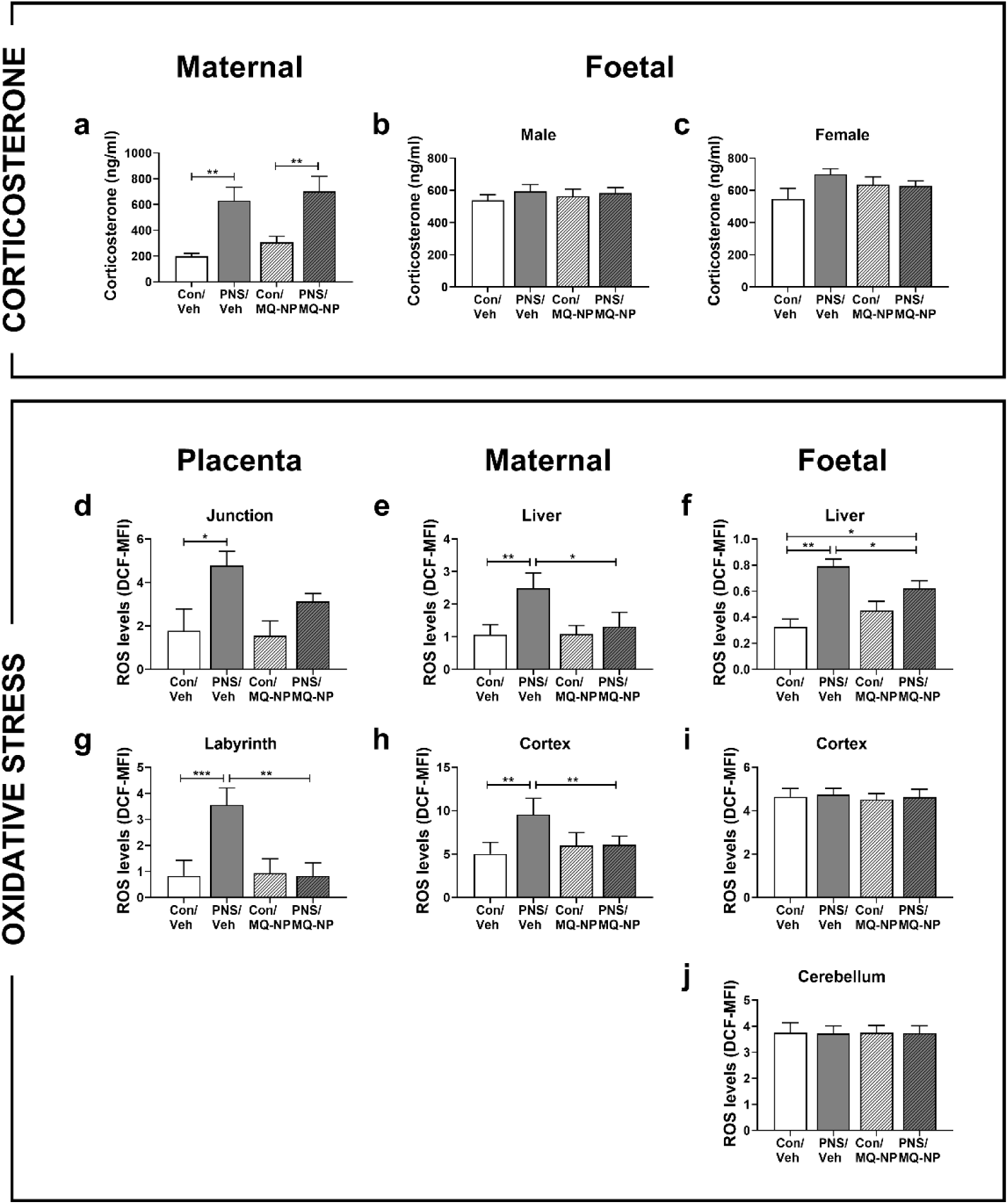
Corticosterone concentrations and oxidative stress levels in the mother and foetuses following prenatal social stress with and without maternal antioxidant treatment. Concentrations of corticosterone were measured in maternal plasma (a), male foetal plasma (b) and female foetal plasma (c) collected on gestational day (GD) 20, from unstressed dams or from dams exposed to 10 min social stress for 5 consecutive days during the last week of pregnancy. Pregnant rats received an intravenous injection of either vehicle or MitoQ-NP prior to the exposure. Levels of dihydrodichlorofluorescein (DCF), a measure of reactive oxygen species (ROS) and a marker of oxidative stress, were quantified at the same time-point in the junctional (d) and labyrinth (g) zones of the placenta, maternal liver (e) and cerebral cortex (h), as well as in the foetal liver (f), cerebral cortex (i) and cerebellum (j). MFI, mean fluorescence intensity. \**p* < 0.05, \*\**p* < 0.01, \*\*\**p* < 0.001 (two-way ANOVA). Group numbers: corticosterone measurements, *n* = 7/group for maternal, male and female samples; DCF levels, *n* = 3/group for maternal, foetal and placental samples.

### 2. Maternal social stress increases oxidative stress in the placenta but not in the foetal brain

There is growing evidence that excessive oxidative stress is associated with adverse outcomes of pregnancy, which in turn are believed to contribute to foetal programming of health and disease in later life^8^. Therefore, we examined whether exposure to psychosocial stress in pregnancy induces oxidative stress in the placenta, and in the maternal and foetal tissues. Following maternal social stress, there was a significant increase in reactive oxygen species in the junctional and labyrinth zones of the placenta (Figure 1d,g), maternal liver and cerebral cortex (Figure 1e,g), and foetal liver (Figure 1f). In contrast, oxidative stress levels in the foetal cerebral cortex and cerebellum were not affected by maternal social stress (Figure 1i,j).

To assess if oxidative stress in the placenta could be prevented or reduced, the pregnant mother was intravenously injected with a nanoparticle-bound mitochondrial antioxidant MitoQ (MitoQ-NP) prior to the social stress exposure. Maternal treatment with MitoQ-NP prevented the increase of oxidative stress in the placental labyrinth (Figure 1g), maternal liver (Figure 1e), maternal cerebral cortex (Figure 1h) and foetal liver (Figure 1f). There was no effect of MitoQ-NP administration on the oxidative stress marker in the foetal brain (Figure 1i,j).

The results suggest that prenatal social stress induces placental oxidative stress, which can be prevented by MitoQ-NP administration to the mother.

### 3. Maternal antioxidant treatment rescues anxiety-like behaviour in male offspring exposed to prenatal social stress

Adult male rats born to mothers exposed to social stress showed increased anxiety-like behaviour, reflected by reduced time spent in the light compartment of the light-dark box (Figure 2a; Supplementary Figure S1) and fewer open arm entries on the elevated plus maze (Figure 2b; Supplementary Figure S1) compared with control males. In contrast, maternal social stress had no effect on anxiety-like behaviour in female rats (Figure 2f,g; Supplementary Figure S1), consistent with our previous findings^4^. Maternal administration of MitoQ-NP prior to social stress exposure prevented the development of anxiety-like behaviour in the male prenatally stressed rats (Figure 2a,b; Supplementary Figure S1) but had no effect in the females (Figure 2f,g; Supplementary Figure S1). CRH-expressing neurons in the central amygdala are especially important in mediating anxious behavioural responses^36,37^. Here, increased anxiety-like behaviour in the male prenatally stressed rats was concomitant with significantly increased *Crh* mRNA expression in the central amygdala (Figure 2c,d), and this effect was prevented by maternal MitoQ-NP treatment in both the adult (Figure 2c,d) and juvenile offspring (P30; Supplementary Figure S1).

**Figure 2.**
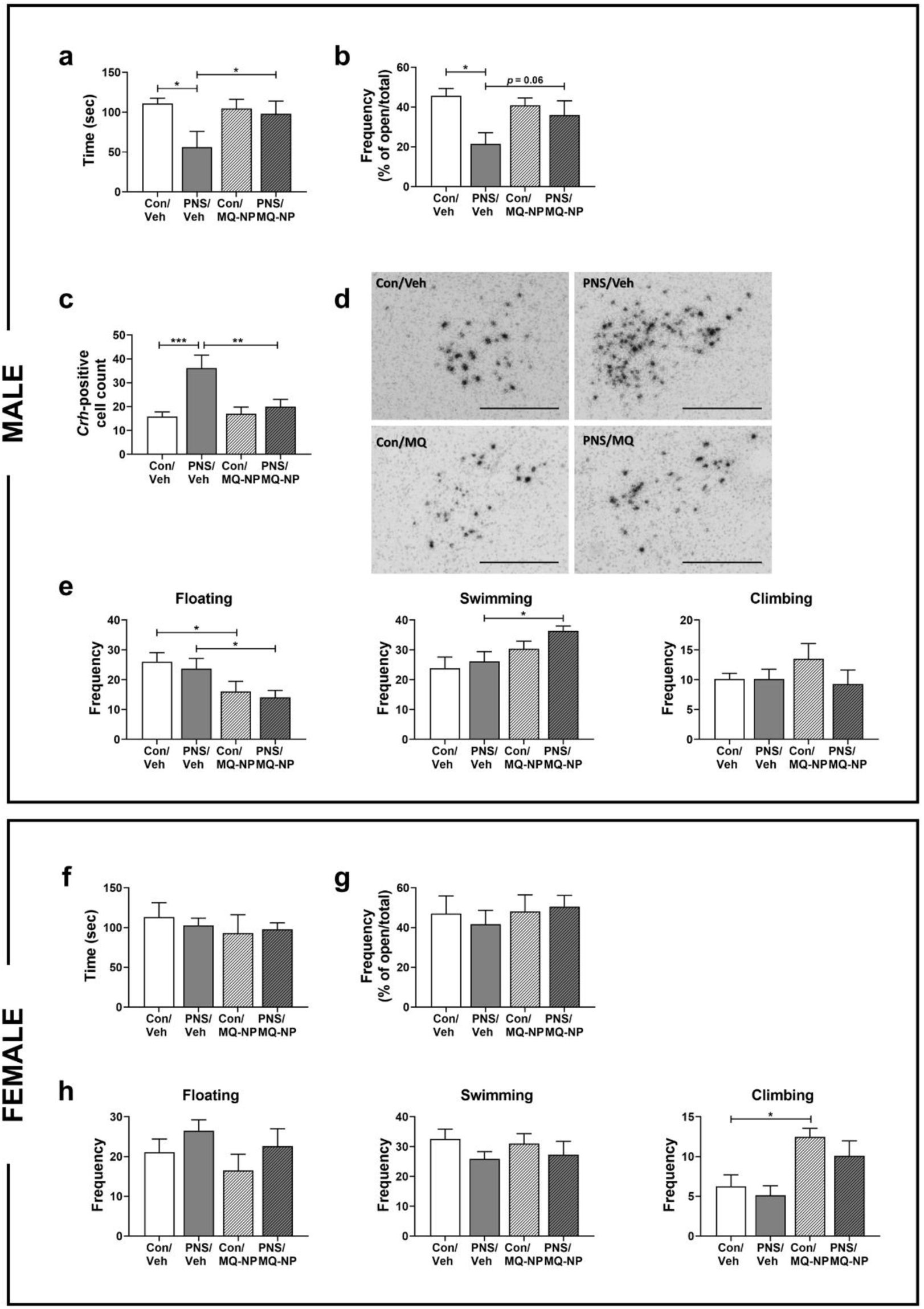
Effects of prenatal social stress and maternal antioxidant treatment on anxiety- and depressive-like behaviour in the adult offspring. The male and female adult offspring of stressed and non-stressed dams, treated with vehicle or MitoQ-NP, were tested for anxiety-like behaviour using the light-dark box (LDB) and the elevated plus maze (EPM), and for depressive-like behaviour using the forced swim test. Time spent in the light compartment of the LDB (a,f) and the percentage of open arm entries on the EPM (b,g) was recorded. In male rats, there was a significant main effect of prenatal stress on the time spent in the light box (F_1,27_ = 4.50, *p* = 0.043; a), with vehicle-treated PNS males spending significantly less time exploring the light box than control males (p = 0.01; a), an effect that was normalised by maternal MitoQ-NP treatment (*p* = 0.027). On the EPM, there was a main effect of prenatal stress status (F_1,26_ = 7.57, *p* = 0.01; b), with prenatally stressed males taking fewer entries into the open arms compared to control males (*p* = 0.0042), and this tended to be normalised by maternal MitoQ-NP treatment (*p* = 0.06). The number of cells positive for *Crh* mRNA expression in the central nucleus of the amygdala was quantified in male adult rats (c). There was a significant main effect of prenatal stress status (F_1,25_ = 12.07, *p* = 0.02), maternal MitoQ-NP treatment (F_1,25_ = 4.99, *p* = 0.035) and also a significant prenatal stress status x maternal treatment interaction (F_1,25_ = 6.79, *p* = 0.015). Representative images of *Crh* mRNA *in situ* hybridisation in the central amygdala of male offspring at postnatal day P30 are shown (d; scale bars = 0.5 mm). Frequency of floating, swimming and climbing behaviour was measured in male and female offspring in the forced swim test (e,h). Two-way ANOVA revealed a significant main effect of MitoQ-NP on floating (F_1,27_ = 9.57, *p* = 0.005) and swimming (F_1,27_ = 7.88, *p* = 0.009; e) in male rats and on climbing in female rats (F_1,27_ = 5.88, *p* = 0.022; h). \**p* < 0.05, \*\**p* < 0.01, \*\*\**p* < 0.001. Group numbers: behavioural experiments, *n* = 8/group for males and females; *Crh* measurements, *n* = 7 for Con/Veh, *n* = 6 for PNS/Veh, *n* = 8 for Con/MQ-NP and PNS/MQ-NP.

There was no significant effect of maternal social stress exposure on depression-like behaviour in either the male or female adult offspring (Figure 2e,h). However, maternal MitoQ-NP administration evidently had an antidepressant action in the forced swim test, independent of prenatal experience (Figure 2e,h), as reflected by reduced passive coping behaviours, (i.e. decreased floating behaviour in male rats), and increased active coping behaviours (i.e. increased swimming in males and increased climbing in females). Neither prenatal stress nor MitoQ-NP treatment affected sucrose preference in either the male or the female offspring (Supplementary Figure S1).

In addition, female offspring exposed to prenatal stress showed reduced social odour memory (Supplementary Figure S1); they were less able to distinguish between novel and familiar social odours compared to controls, consistent with previous findings^5^. This effect was not significantly altered by maternal MitoQ-NP treatment (Supplementary Figure S1).

In summary, administration of an antioxidant to the mother prior to social stress exposure prevented stress-induced changes in anxiety-like behaviour in adult male offspring and seemingly had anti-depressant effects in both sexes, regardless of prenatal experience.

### 4. Prenatal social stress leads to differential expression of serotonin and noradrenaline-related genes in the foetal frontal cortex

To establish the impact of prenatal stress exposure on central gene expression, RNA sequencing was performed. Following prenatal stress, 143 genes were found to be differentially expressed in the foetal frontal cortex most of which were downregulated (12 upregulated, 131 downregulated) (Supplementary Data S1). Enriched pathways corresponded to serotonin and adrenaline/noradrenaline biosynthesis (Figure 3a). MitoQ-NP treatment reduced the prenatal stress-induced up or downregulation of cortical genes, as demonstrated by a negative correlation between fold change differences in gene expression induced by prenatal stress (compared to non-stressed controls) and fold change differences induced by prenatal stress with MitoQ-NP treatment (compared to prenatally stressed rats that had received saline treatment) (Spearman correlation = −0.798, p < 2.2E-16). Genes related to serotonin and noradrenaline biosynthesis, transport or differentiation were generally found to be downregulated and maternal MitoQ-NP treatment significantly prevented the downregulation of three of these genes in the foetal brain (Figure 3b). Western blot analysis of the protein products in foetal frontal cortices confirmed the significant downregulation of tyrosine hydroxylase in offspring exposed to prenatal stress compared to control offspring (Figure 3c). In contrast, in the juvenile offspring (P30) tyrosine hydroxylase expression was significantly greater in the frontal cortex, following prenatal stress (Figure 3d). Protein levels of dopa decarboxylase tended to be reduced in the cortex of foetal offspring in response to prenatal stress (Figure 3e), however this effect was not significant (*p* = 0.095). In the juvenile offspring, prenatal stress exposure did not affect dopa decarboxylase levels in the frontal cortex (Figure 3f). As observed by RNA sequencing, maternal MitoQ-NP treatment did not significantly affect the levels of tyrosine hydroxylase or dopa decarboxylase in the frontal cortex of the foetuses (Figure 3c,e), or the juveniles (Figure 3d,f).

**Figure 3.**
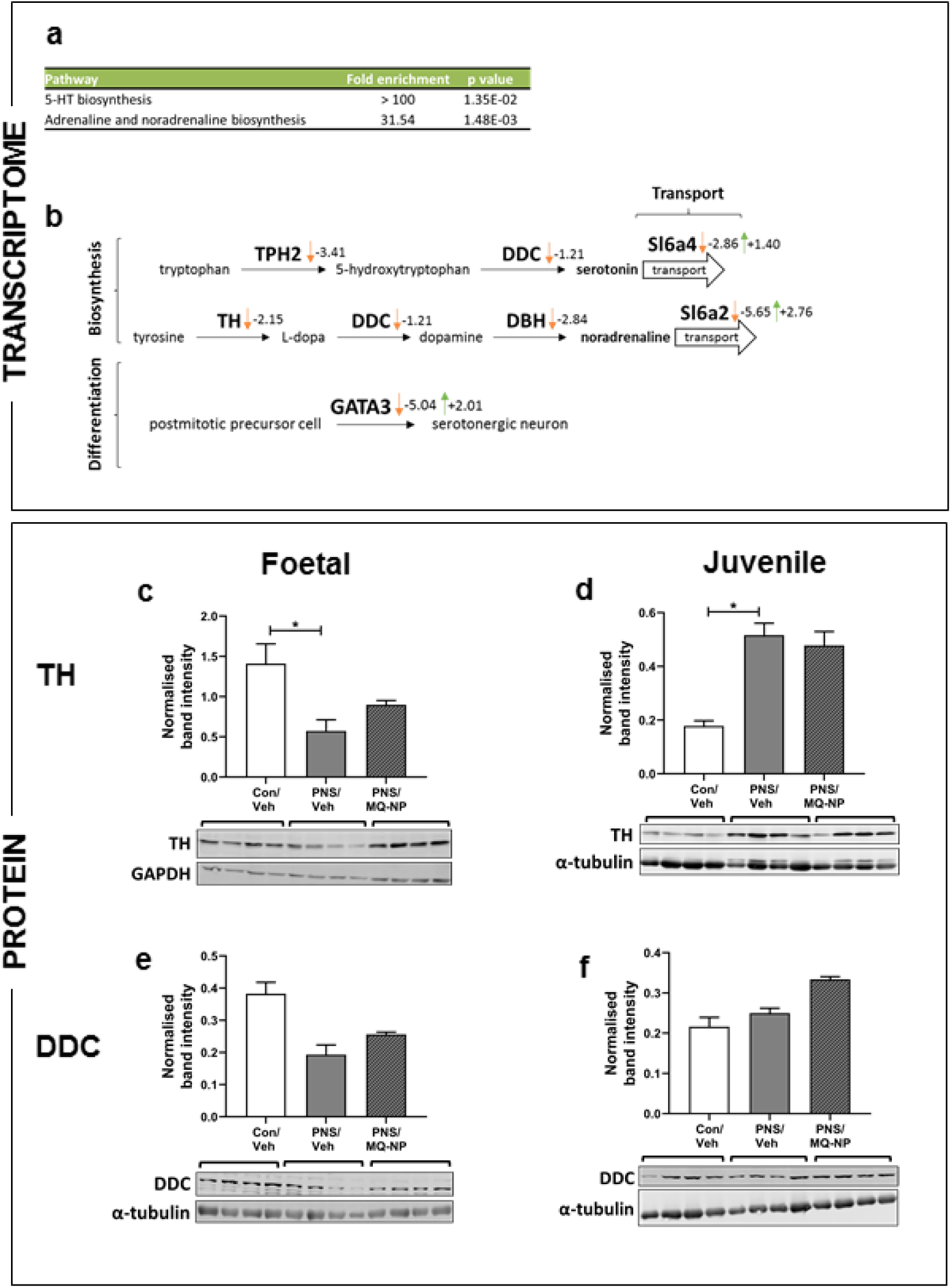
Gene and protein expression changes in the offspring brain in response to maternal social stress and antioxidant treatment. RNA (a-b) and protein levels (c-f) were analysed in cerebral cortex tissue collected from foetal (a-c,e) and P30 juvenile (d,f) offspring of non-stressed mothers and those exposed to social stress with either vehicle or MitoQ-NP pretreatment. Significant differentially expressed genes, identified by RNA sequencing, from prenatally stressed rats compared to control rats were subjected to pathway analysis (a). Differentially expressed genes involved in serotonin and noradrenaline-related processes are shown with orange arrows indicating differential expression under prenatal stress conditions and green arrows representing differential expression following prenatal stress combined with maternal MitoQ-NP treatment. Values represent log-2 fold changes (b). Relative protein levels of tyrosine hydroxylase (TH; c,d) and dopa decarboxylase (DDC; e,f) were quantified by Western blot. One-way ANOVA showed a significant difference between experimental groups of tyrosine hydroxylase expression in the frontal cortex of foetal (F_2,11_ = 5.07, *p* = 0.034) and juvenile offspring (F_2,11_ = 6.53, *p* = 0.018). \**p* < 0.05. Group numbers: *n* = 4/group for foetal and juvenile tissue.

The results indicate that prenatal stress alters the expression of serotonin and noradrenaline-related genes in the foetal brain; these changes were transient and recovered by P30. Nonetheless, some of these changes could be prevented by administration of MitoQ-NP to the mother.

### 5. Maternal antioxidant treatment prevents neurological changes in the adult offspring exposed to prenatal social stress

Exposure to prenatal social stress induced several neuroanatomical and neurochemical changes in the adult offspring’s brain. In the male adult offspring, prenatal social stress resulted in a significant reduction in dendrite length in the CA3 region of the hippocampus, basolateral amygdala (Figure 4a), auditory cortex, somatosensory cortex and thalamic reticular nucleus (Supplementary Figure S2). In contrast, in female offspring a reduction in dendritic length was only observed in the auditory cortex (Supplementary Figure S2), but not in any of the other brain regions examined (Figure 4e). Maternal administration of MitoQ-NP significantly prevented the prenatal stress-induced reduction in dendritic length in the hippocampal CA3 region (Figure 4a), somatosensory cortex and thalamic reticular nucleus of the male offspring (Supplementary Figure S2), but the same treatment had no effect in the female offspring (Figure 4e; Supplementary Figure S2). Process lengths of tyrosine hydroxylase-positive (TH+) neurons were significantly reduced in the basolateral amygdala of both male and female offspring, and this effect was prevented by maternal treatment with MitoQ-NP (Figure 4b,f). TH+ process lengths in the hippocampus (Figure 4b,f) and cerebral cortex were not significantly affected by prenatal stress or drug treatment in either sex (Supplementary Figure S2).

**Figure 4.**
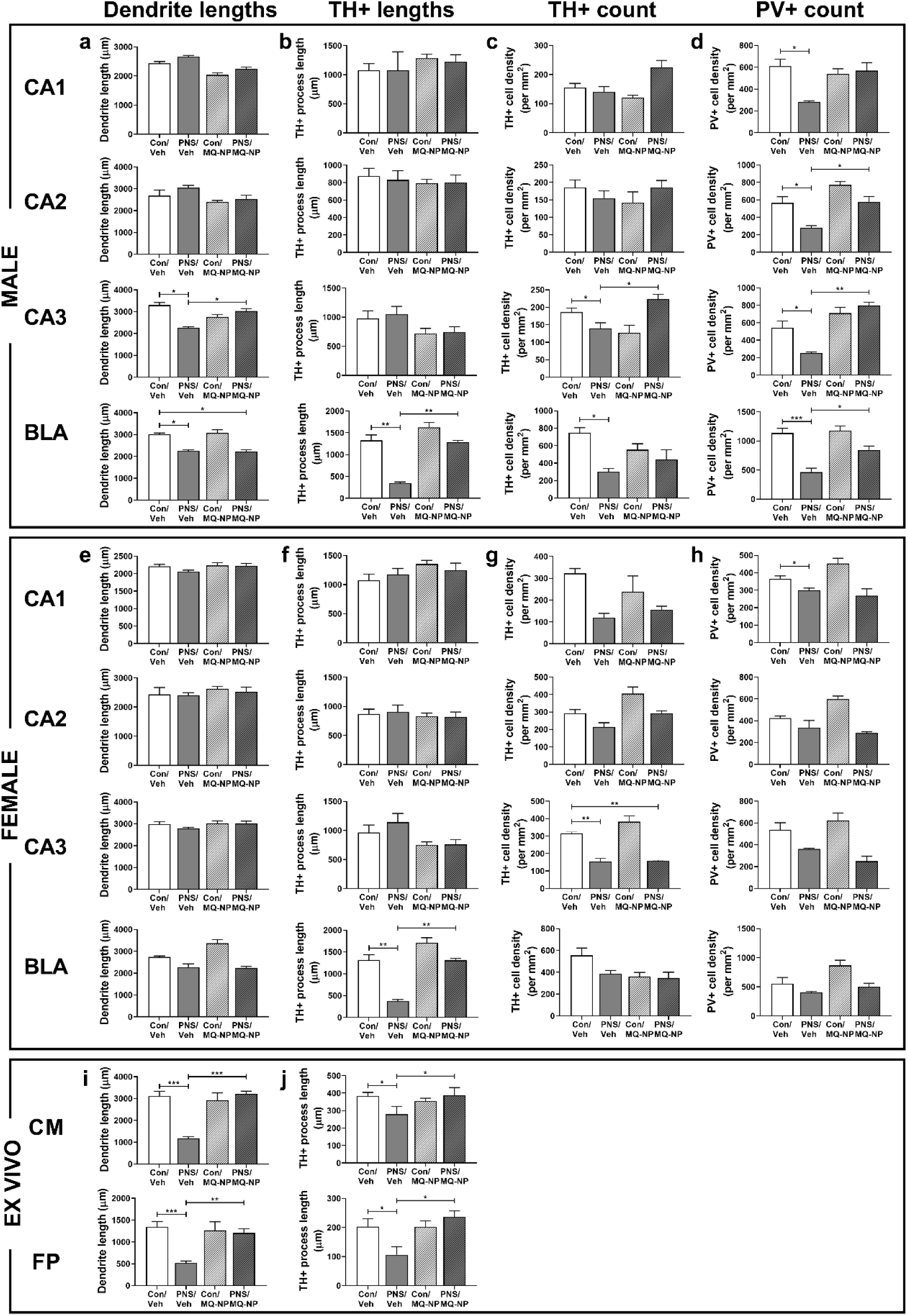
Neuroanatomical effects of maternal social stress and antioxidant treatment in juvenile offspring *in vivo* and on cortical cultures *in vitro*. Brain tissue from juvenile rats, born to stressed or non-stressed mothers administered either vehicle or MitoQ-NP during pregnancy, were processed using immunohistochemistry (a-h). Dendrite lengths (a,e), lengths of TH+ cell processes (b,f), TH+ cell numbers (c,g) and PV+ cell numbers (d,h) were assessed in the CA1, CA2, CA3 hippocampal subregions and in the basolateral amygdala (BLA) in male and female offspring. Foetal plasma (FP) and culture medium conditioned by placentae collected on gestational day 20 from dams of the four different treatment groups were applied to embryonic cortical culture (i,j). Dendrite lengths (i) and TH+ process lengths (j) are shown. \**p* < 0.05, \*\**p* < 0.01, \*\*\**p* < 0.001. Group numbers: *n* = 6/group for males and females.

Prenatal stress did not alter the overall density of neurons in male or female offspring in any of the regions examined (Supplementary Figure S3). Analysis of the number of parvalbumin-positive (PV+) and TH+ neurons, revealed a loss of immunopositive neurons in the hippocampal subfields, cortex, amygdala and reticular nucleus in male and female prenatally stressed offspring (Figure 4a,d,g,h; Supplementary Figure S2). Specifically, the number of TH+ neurons was significantly lower in the CA3 and the basolateral amygdala of male offspring, as well as in the CA3 and thalamic reticular nucleus of female offspring, compared with controls (Figure 4c,f; Supplementary Figure S2). PV+ cell density was significantly lower in all examined subfields of the hippocampus and in the cerebral cortex, along with the basolateral amygdala and thalamic reticular nucleus in male prenatally stressed offspring, compared with controls (Figure 4d; Supplementary Figure S2), while in the female prenatally stressed offspring significantly lower numbers of PV+ neurons were observed in CA1, auditory cortex, retrosplenial cortex and thalamic reticular nucleus (Figure 4h; Supplementary Figure S2). Maternal MitoQ-NP treatment prevented the effects of prenatal stress on PV+ neurons in most brain regions (Figure 4d,h; Supplementary Figure S2), but only rescued the reduced TH+ cell density in the hippocampal CA3 region of male offspring (Figure 4c,f; Supplementary Figure S2).

Immunoreactivity for the GABA receptor subunits Aα1, Aα2 and B1 was significantly lower in the CA1 and CA2 hippocampal subfields, basolateral amygdala, thalamic reticular nucleus, and auditory and somatosensory cortex of prenatally stressed male offspring, compared to male control offspring (Figure 5a-c; Supplementary Figure S4). In female prenatally stressed offspring, GABA Aα1 and Aα2 levels were reduced in the CA1 and somatosensory cortex (Figure 5e,f), while GABA B2 immunoreactivity was significantly reduced in CA1, CA2, basolateral amygdala and all examined subregions of the cerebral cortex (Figure 5g; Supplementary Figure S4). Maternal administration of MitoQ-NP prevented the prenatal stress-induced reduction in GABA Aα1 in the female CA1 (Figure 5e), GABA Aα2 in the male and female hippocampus and in the male basolateral amygdala (Figure 5b,f), and GABA B1 in male CA1 and basolateral amygdala, along with male and female cortical subregions (Figure 5c; Supplementary Figure S4). Expression of the glutamate receptor subunit GluN1 in male prenatally stressed offspring was significantly lower in the CA2, thalamic reticular nucleus and somatosensory cortex, compared to control males; while GluN1 immunoreactivity was significantly greater in the auditory cortex (Figure 4d; Supplementary Figure S4). In female prenatally stressed offspring, lower expression of GluN1 in the CA2, CA3, basolateral amygdala and all cortical subregions was observed, compared with control females (Figure 5h; Supplementary Figure S4). MitoQ-NP treatment was effective in preventing the prenatal-stress induced reduction in GluN1 subunits in the somatosensory cortex and thalamic reticular nucleus in male rats (Supplementary Figure S4), as well as in the basolateral amygdala, auditory and somatosensory cortex of female rats (Figure 5h; Supplementary Figure S4).

**Figure 5.**
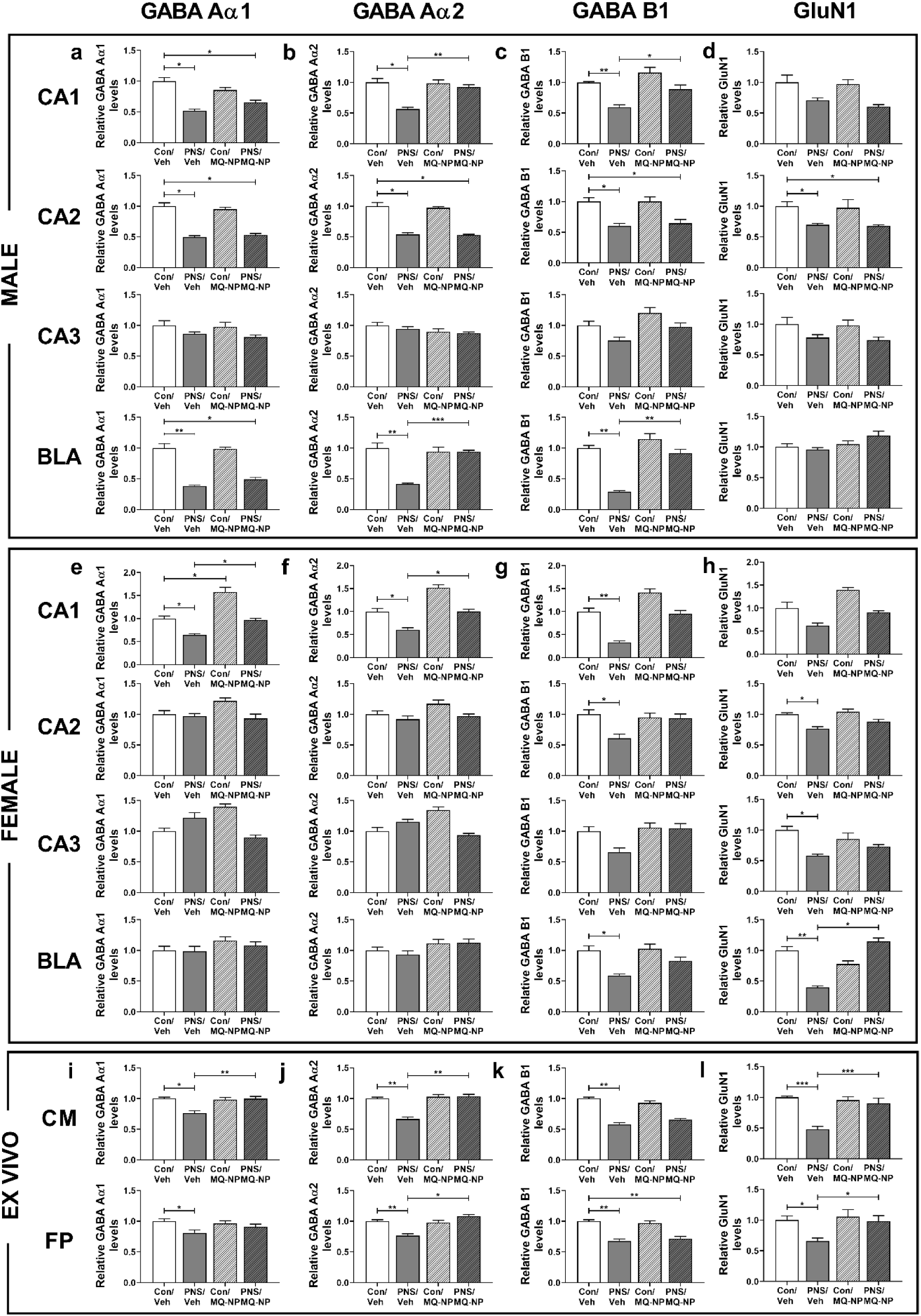
Neurochemical effects of maternal social stress and antioxidant treatment in juvenile offspring *in vivo* and on cortical cultures *in vitro*. Immunoreactivity for GABA and glutamate receptor subunits was assessed in the brains of juvenile rats, born to mothers either exposed to social stress or undisturbed during pregnancy, and pre-treated with either vehicle or MitoQ-NP. GABA Aα1 (a,e), GABA Aα2 (b,f), GABA B1 (c,g) and GluN1 (d,h) subunit expression was quantified in the hippocampal subregions CA1, CA2, CA3 and in the basolateral amygdala (BLA) in both male and female offspring. Embryonic cortical cultures were exposed to foetal plasma or culture medium conditioned by placentae, collected at gestational day 20 from pregnancies exposed to social stress and/or MitoQ-NP treatment (i-l). Expression of GABA Aα1 (i), GABA Aα2 (j), GABA B1 (k) and GluN1 (l) subunits was quantified in cortical cell cultures. \**p* < 0.05, \*\**p* < 0.01, \*\*\**p* < 0.001. Group numbers are: *n* = 6/group for males and females.

Taken together, the results indicate that prenatal social stress exposure causes long-term neuroanatomical changes in the juvenile offspring brain, and support a role for dysregulated GABAergic and glutamatergic signalling in prenatally stressed offspring. Many of these neuroanatomical and neurochemical changes are prevented by maternal MitoQ-NP administration prior to the stress exposure.

### 6. Exposing cortical cultures to foetal plasma or placenta-conditioned medium from stressed mothers mimics the neurological effects of prenatal stress

Due to the observed lack of increased corticosterone in the foetal circulation in response to maternal social stress, we hypothesised that the placenta, as the interface between mother and foetus, might secrete factors into the foetal circulation that alter foetal brain development. To test this hypothesis, neuronal cell cultures from dissociated cortical tissue were exposed to foetal plasma collected from pregnant rats that were either undisturbed or exposed to social stress, with and without MitoQ-NP pretreatment. Additionally, placental tissue from the same pregnancies was cultured *ex vivo* and the resulting placenta-conditioned medium was applied to cortical cultures.

A significant reduction in dendritic length was observed in cortical cultures following exposure to placenta-conditioned medium or foetal plasma collected from prenatally stressed rats (Figure 4i), similar to the changes observed in the brains of the male offspring (Figure 4a). Maternal MitoQ-NP treatment prevented these stress-induced effects on dendrite length (Figure 4i). Exposure to conditioned medium or foetal plasma from stressed rats, resulted in shorter TH+ process lengths (Figure 4j), in line with our observations in the basolateral amygdala of prenatally stressed offspring (Figure 4b,f). This effect of maternal stress on TH+ process length was not observed in the cortical cultures exposed to foetal plasma or placenta-conditioned medium collected from stressed mothers treated with MitoQ-NP (Figure 4j). As observed *in vivo*, the total density of neurons was not altered in cortical cultures exposed to conditioned medium or foetal plasma from any treatment groups (Supplementary Figure S3). Because of the paucity of TH+ and PV+ cells in cortical cultures, no meaningful counts for these cell types could be obtained.

Immunoreactivity for GABA receptor subunits Aα1, Aα2 and B1, as well as glutamate receptor subunit GluN1 was significantly lower in cortical cultures exposed to either conditioned medium or foetal plasma from stressed rats (Figure 5i-l), replicating the loss of immunoreactivity for these receptors seen across multiple brain regions in both the male and female prenatally stressed offspring (Figure 5a-h; Supplementary Figure S4). Maternal MitoQ-NP treatment was successful in preventing the reduced levels of GABA Aα1, GABA Aα2 and GluN1 in cortical cultures exposed to conditioned medium (Figure 5i,j,l) and levels of GABA Aα2 and GluN1 in cultures exposed to foetal plasma (Figure 5j,l).

In summary, placenta-conditioned medium and foetal plasma induced morphological and neurochemical changes in cortical neurons *in vitro*, similar to those observed in distinct brain regions of juvenile offspring exposed to prenatal stress, suggesting in response to stress the placenta may secrete an as yet unidentified factor which accesses the foetal blood. Prenatal treatment of the mother with MitoQ-NP prevented many of these changes.

### 7. microRNAs are differentially secreted from the placenta in response to prenatal social stress

The evidence suggests the presence of factors in the culture medium conditioned by placental tissue and in the foetal plasma in response to maternal stress, that induce similar changes in dissociated cortical neurons to those changes observed in the brains of prenatally stressed juvenile offspring. Hence we conducted an initial search for potential mediators of these changes.

microRNAs are small post-transcriptional regulators that pay a role in neurodevelopment and in neurodevelopmental disorders, including anxiety and depression^38,39^. We detected differentially abundant microRNAs in response to prenatal stress in both conditioned medium and foetal plasma (Figure 6a; Supplementary Data S2). Ten microRNAs were similarly up or downregulated in both placenta-conditioned medium and foetal plasma from stressed rats and may therefore be of particular interest for further study. microRNAs significantly altered by prenatal stress consisted of a higher proportion of hypoxamiRs, hypoxia-regulated microRNAs and this enrichment was significant for microRNAs in conditioned medium (Figure 6b). Gene ontology analysis of predicted targets of the significantly altered microRNAs showed enrichment in similar biological processes for conditioned medium and foetal plasma. microRNA targets were enriched for nervous system development and signalling pathways (Figure 6c). Maternal MitoQ-NP administration prevented abnormal changes to 12 of the significant microRNAs in conditioned medium and 4 microRNAs in foetal plasma (Supplementary Data S2). miR-455 was found to be significantly upregulated by prenatal stress in both conditioned medium and foetal plasma, as well as significantly reduced to control levels by MitoQ-NP treatment (Supplementary Data S2).

**Figure 6.**
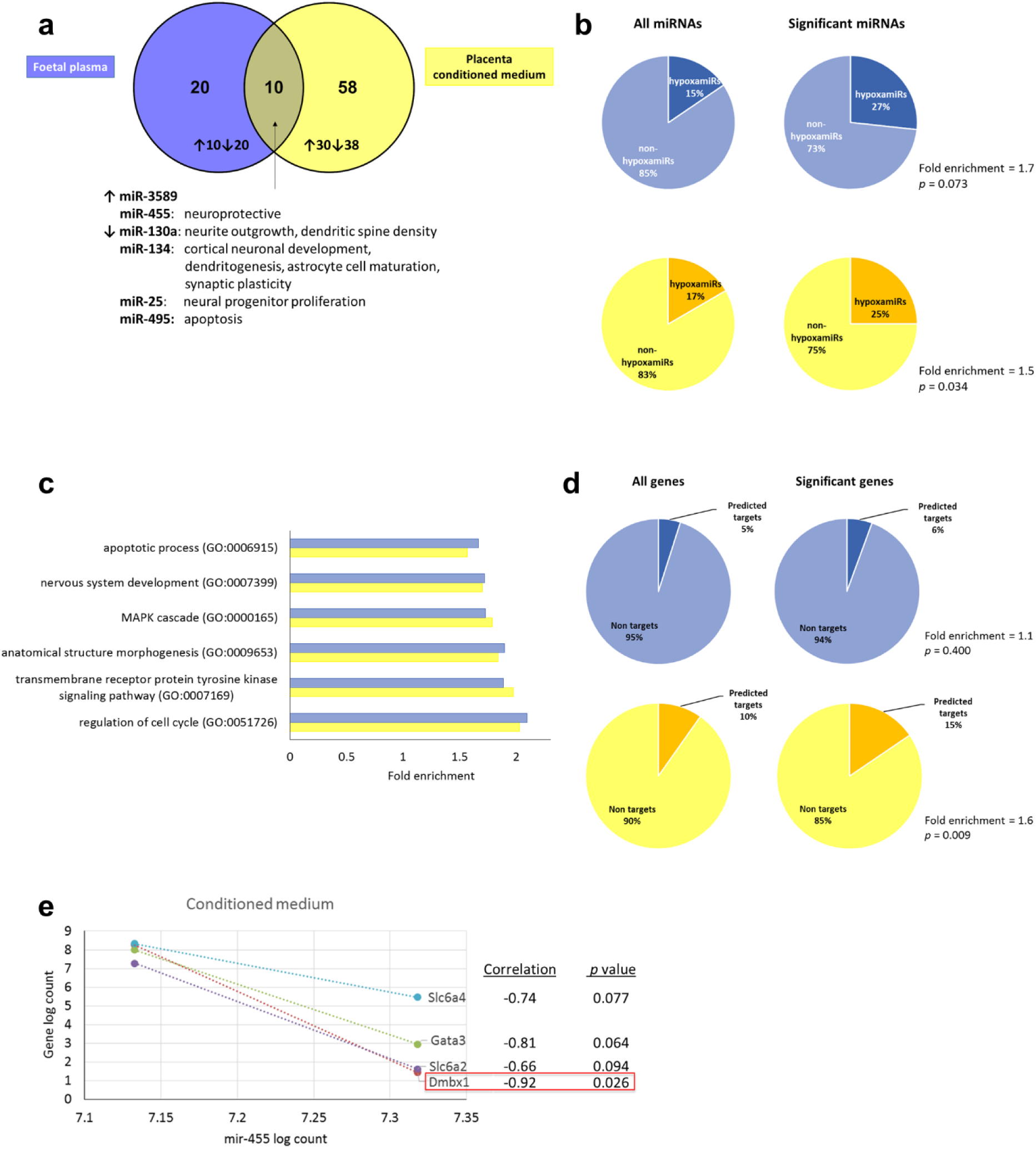
Circulating microRNAs in foetal plasma and placenta-conditioned medium following prenatal social stress. Pooled foetal plasma (blue), along with placenta-conditioned culture medium (yellow), collected from pregnant rats exposed to prenatal social stress and treated with vehicle, was assessed for levels of microRNAs. Differentially abundant microRNAs following prenatal stress were compared between foetal plasma and placental conditioned medium (a). Significant differentially abundant microRNAs were enriched for hypoxamiRs (b), both in foetal plasma (top) and placenta conditioned medium (bottom). Gene ontology analysis was performed on the predicted targets of differentially abundant microRNAs (c). Predicted targets of differentially abundant microRNAs were analysed for enrichment among differentially expressed genes in cortex of foetuses from the same pregnancies (d). Correlation between levels of genes *Dmbx1*, *Gata3*, *Slc6a2* and *Slc6a4* in the foetal cortex with levels of miR-455 in placental conditioned medium is shown (e). The red box signifies a significant correlation (*p* < 0.05). Group numbers: *n* = 3/group for plasma and conditioned medium.

Predicted targets of those microRNAs that showed significantly different abundance in placenta-conditioned medium were specifically enriched among the differentially expressed genes in the foetal cortex detected by RNA-sequencing (Figure 6d). Correlation analysis of miR-455 with the four differentially expressed genes that had been shown to be affected by both prenatal stress and MitoQ-NP treatment (*Dmbx1*, *Gata3*, *Slc6a2* and *Slc6a4*), suggested a negative correlation between gene expression in the foetal frontal cortex and levels of mir-455 secreted from the placenta, i.e. transcript levels of the genes were downregulated by prenatal stress compared to non-stressed rats, while miR-455 levels in conditioned medium were upregulated (Figure 6e). This correlation was significant for the transcription factor *Dmbx1*.

The results suggest that the pattern of microRNAs released from the placenta is altered following maternal social stress. Some of these changes can be prevented by maternal MitoQ-NP treatment and may be correlated with gene expression changes in the foetal brain.

## Discussion

Psychosocial stress during pregnancy has been associated with an increased risk of psychiatric and neurodevelopmental disorders in the offspring^1–3^. In the present study, we observed long-term neuroanatomical, neurochemical, behavioural and gene expression changes in the offspring of rats that experienced social stress during pregnancy.

The mechanism by which the effects of maternal social stress are conveyed to the foetus is unclear. Previous studies have focussed, almost exclusively, on a role for maternal glucocorticoids^7,8^. In the present study however, despite a marked stress-induced increase in corticosterone secretion in the mother, we did not observe any such increase in corticosterone concentrations in the foetal circulation at gestational day 20, indicating that maternal corticosterone is unlikely to directly program the foetal brain. Whether maternal glucocorticoids contribute to foetal programming indirectly by initiating a cascade of downstream events, for example by interacting at the placenta, remains to be tested.

Previous research has suggested that the placenta, as the interface between mother and foetus, plays a key role in neurodevelopment and in foetal programming of psychological disorders that result from adverse events during pregnancy^31,40,41^. In keeping with this concept, the results presented here demonstrate for the first time that maternal exposure to social stress during pregnancy induces oxidative stress in the placenta, highlighting the potential significance of the placenta in mediating the effects of social stress on the offspring. This observation is in line with previous studies showing that placental oxidative stress is increased in response to other maternal insults during pregnancy, such as hypoxia^31^.

Maternal administration of a nanoparticle-bound antioxidant, MitoQ-NP, prior to the chronic social stress exposure was able to reduce placental and maternal oxidative stress, as seen previously in pregnant rats exposed to hypoxia^31^. Moreover, a single MitoQ-NP injection to the mother was able to prevent anxiety-like behaviour in male offspring exposed to prenatal stress and had an overall antidepressant effect in males and females, regardless of maternal stress exposure. This observation further suggests a potential role for oxidative stress in the placenta in the foetal programming of long-term disease and suggests that these long-term effects can be prevented through treatment of the mother during pregnancy. The advantage of using nanoparticle-bound MitoQ lies in the observation that it does not enter the foetal brain, as previously shown^31^, thereby reducing the risk of potential side effects to the developing foetus. MitoQ is a mitochondrial targeted antioxidant that is repeatedly recycled back to its active form by the respiratory chain after fulfilling its antioxidant role^42,43^ and can thus produce long-lasting effects. In light of the evident need for preventative treatment, the observation that maternal MitoQ-NP treatment prevents behavioural changes in the offspring in later life warrants further assessment.

A comparison was made of the neurological changes between male and female adult offspring in view of the specific vulnerability of the male offspring to increased anxiety-like behaviour. Particularly marked neurological changes were observed in the basolateral amygdala, hippocampus and the thalamic reticular nucleus and there was a general tendency for male offspring to show more pronounced neurological changes compared to females, consistent with other studies that report sex-specific foetal programming in response to maternal insults during pregnancy^44–47^. For example, here the finding that GABA_A_ receptor subunit expression was lower in the BLA of prenatally stressed males, but not females, may be important particularly given only the males displayed an anxious phenotype and changes in both receptor expression and anxiety behaviour were prevented by maternal antioxidant treatment. Indeed, decreased GABAergic transmission in the BLA is strongly linked to heightened anxiety behaviour in rodents and reduced GABA_A_ receptor binding has been reported in the brains of patients with generalised anxiety disorder^32,48–50^. Moreover, the observed reduction in the number of PV+ neurons was more widespread in prenatally stressed males than in females, which may be particularly relevant to anxiety behaviour, as impaired inhibitory drive from PV+ interneurons is consistently reported in animal models of psychiatric disorders and an abundance of evidence supports disrupted inhibitory control in several psychiatric disorders^33,34^.

Some prenatal stress-induced neurological changes were only observed in the male offspring. These included a shortening of the dendrites in the basolateral amygdala, hippocampal CA3 region and thalamic reticular nucleus; a decrease in PV+ cells in basolateral amygdala and CA3, and a reduction in TH+ cells in the basolateral amygdala. The basolateral amygdala, hippocampus and thalamus are thought to play key roles regulating anxiety-related behavioural responses^35,51,52^. In particular, a role for PV+ neurons in the basolateral amygdala behaviours has been reported in the regulation of anxiety-related behaviours^53–55^. Furthermore, dendrite remodelling in hippocampus and amygdala has been observed previously in response to chronic stress^56,57^. The selective impact on the CA3 region supports the hypothesis that the hippocampus is not a unitary structure but consists of subfields with specific functions. Through region-specific knockouts the CA3 has been shown to play an essential role in anxiety-like behaviours, as opposed to fear responses^58^.

Why these brain regions are particularly susceptible to stress-induced changes in male, but not female rats is unclear. Differences in sex steroids may render certain brain regions more vulnerable to prenatal stress in males versus females; however, differences in gene expression or epigenetic profiles are also likely to play a role^59,60^. In the basolateral amygdala differential expression of CRH receptor subtypes has been observed in male rats exposed to prenatal stress, but not in females^61^. Furthermore, sex-specific differences in the placenta, specifically differences in gene expression^20,22,26,62,63^, epigenetic profiles^24,28^, and metabolism^64^, could render male and female placentae differentially responsive to maternal insults^45,65^.

Application of culture medium conditioned by placentae, or foetal plasma, collected from mothers exposed to social stress to neuronal cultures, replicated many of the neurological effects observed in the adult offspring, such as reductions in dendrite length, TH+ process lengths and levels of GABA and glutamate receptor subunits. These effects were prevented when using conditioned medium or plasma from mothers who had been treated with MitoQ-NP. Due to the low volume of foetal plasma that could be collected from each foetus, samples were pooled across litters, meaning that no comparisons could be made between males and females. However, the above suggests that in response to maternal social stress the placenta secretes molecules into the foetal circulation that can alter the foetal brain. We have previously proposed that the placental barrier may respond to a variety of physiological insults by secreting factors that can damage cells distal to the barrier, both *in vitro* and *in vivo*, thereby having the potential to play a role in foetal development^31,40,41,66,67^. The current work further supports the hypothesis that factors released from the placenta may mediate the effects of maternal insults during pregnancy on the developing foetal brain.

Preliminary exploration of potential factors that were secreted from the placenta showed that microRNAs were differentially abundant both in placenta conditioned medium and foetal plasma from pregnant rats exposed to social stress, compared to unstressed rats. Some of these changes were prevented by maternal administration of MitoQ-NP. Furthermore, a potential correlation between microRNA released from the placenta and gene expression changes in the foetal brain was observed. Extracellular microRNAs can act as long-distance messengers. They can be taken up by cells, including neurons, where they fulfil their regulatory function^68^. Furthermore, microRNAs have the potential to cross the blood-brain barrier and enter the brain, when encapsulated within extracellular vesicles^69^. We hypothesise that following chronic maternal social stress, extracellular microRNAs released into the foetal circulation from the oxidatively stressed placenta may cross the blood-brain barrier, where they can be taken up by neurons and transiently affect the expression of crucial developmental genes. In the foetal cortex of offspring exposed to prenatal stress, we observed a reduction in the expression of genes associated with serotonin and noradrenaline biosynthesis and transport. However in the prenatally stressed juveniles, protein levels in the frontal cortex had returned to levels equivalent to those observed in non-stressed offspring. The serotonin system plays a critical role during neurodevelopment, in particular in the development of the limbic system (including the amygdala, hippocampus and prefrontal cortex), which is important for processing emotional behaviours^70^. Therefore, this transient dysregulation of gene expression in prenatally stressed foetal brains could temporarily affect crucial neurodevelopmental processes at a period of high vulnerability, in line with the concept of the foetal origins of adulthood disease^71,72^, potentially leading to long-term alterations in the offspring brains. Sex-specific differences at any level of this process (e.g. differences in the levels of secreted factors, uptake into the brain, or a specific vulnerability of particular brain regions) may lead to differences in neurological and behavioural phenotypes in adulthood in males and female offspring, although this remains to be tested.

### Conclusion

The prenatal period has been shown to be critical for healthy development. However, very little in the way of preventative treatment exists for cases when the prenatal environment is sub-optimal (e.g. during periods of chronic maternal stress). The present study highlights for the first time the importance of oxidative stress in the placenta – and factors released from the placenta that likely access the foetal circulation – in mediating the effects of maternal psychosocial stress exposure on the offspring’s brain and behaviour. While further assessment of maternal antioxidant treatment in pregnancy will be required, especially in light of the observed differences in male and female offspring, the present study highlights the potential for drugs such as MitoQ-NP to be used as preventative treatment with long-term implications for mental health.

## Methods

### Experimental design and animals

All animal procedures were conducted in accordance with the UK Animals (Scientific Procedures) Act 1986 and the European Directive (2010/63/EU) and with institutional approval from the Roslin Institute Animal Welfare and Ethical Review Body at the University of Edinburgh.

Female Sprague-Dawley rats (initial body weight: 240–250g; Charles River, Margate, Kent, UK) were maintained on *ad libitum* standard rat chow (Teklan 2014 diet, Harlan Laboratories, UK; supplemented 1:1 with Teklan 2019 diet throughout pregnancy and lactation) and filtered tap water under a 12:12-h light-dark cycle (lights on at 07:00h). Following ≥ 7 d acclimatisation to the facility, female rats were housed overnight with a sexually experienced male. Pregnancy was confirmed by the presence of a semen plug in the breeding cage the following morning and this was designated day 1 of pregnancy. Rats were group housed (4-6/cage) in individually ventilated cages until gestational day (GD) 16 and individually thereafter.

### Maternal social stress paradigm and drug treatment

On GD16, pregnant rats were weighed and 4% (w/w) lidocaine cream was applied topically to the tail prior to administration of either 0.9% sterile saline (vehicle; 0.34 ml/kg) or 125 µM nanoparticle-bound mitoquinone (MitoQ-NP; 0.34 ml/kg) i.v. into the tail vein. Immediately after the injection, each rat was transferred into a clean cage. Approximately 1 h later, half of the vehicle and MitoQ-NP-treated rats were exposed to 10 min social stress, while the remainder served as non-stressed controls and remained undisturbed in their home cage. Social stress was induced using a modified resident-intruder paradigm as previously described^4^. Briefly, pregnant rats (‘intruder’) were placed in the home cage of an unfamiliar aggressive lactating rat (‘resident’ between days 1-7 of lactation) for 10 min/day for 5 consecutive days from GD16-20. A different resident rat was used on each day. The number of attacks and latency to the first attack were scored manually. All rats were weighed daily from GD 16-20.

Two sets of in *vivo* experiments were performed:

#### Experiment 1: Effects of prenatal stress on the mother, foetus and placenta at GD20

Pregnant rats underwent drug treatment and social stress exposure (as above). Immediately after the final social stress exposure on GD20, pregnant rats and foetuses were killed by conscious decapitation. Maternal trunk blood was collected into chilled tubes containing 5% (w/v) EDTA. Foetal trunk blood was collected into EDTA-coated microvette tubes (Sarstedt, Germany), pooled by sex and litter. In both cases, plasma was separated by centrifugation at 4°C and then frozen on dry ice and stored at −20°C until analysis of corticosterone and microRNA levels along with application to cortical cultures (see below). Placental explants (n = 3/litter) were collected and cultured as described below to generate conditioned media. Additional placentae, together with maternal and foetal samples of brain and liver were also collected and frozen on dry ice before storage at −80°C for oxidative stress, transcriptome and protein analyses (see below).

#### Experiment 2: Effects of prenatal stress on offspring brain and behaviour

An additional cohort of pregnant rats underwent the same drug treatment and social stress exposure (as above) but were allowed to give birth (typically on GD22.5-23). Pups remained with their mothers until weaning on postnatal day (P) 23. Litter weights were measured on the day of birth, on P8 and at weaning (P23). Following weaning, offspring were housed in same sex groups by litter (4-6 females, 3-5 males) and were left undisturbed except for handling during routine husbandry. At P30, a cohort of rats (n = 6-7 males and n = 6-7 females per treatment group each from separate litters) were deeply anaesthetised with 3% isoflurane in 1000 ml/min O_2_, before cardiac perfusion fixation. Following fixation, brains were removed and transferred into 4% (w/v) paraformaldehyde in 15% (w/v) sucrose solution and stored overnight at 4°C, after which time they were transferred to 30% (w/v) sucrose solution at 4°C for ca. 48 h, after which time they were frozen on dry ice and stored at −80°C until processing for immunohistochemistry (see below). Another cohort of male offspring (n = 5/treatment group each from different dams) were killed by CO_2_ overdose and the prefrontal cortex was collected for protein analysis (see below). The remaining littermates were group housed (by litter and sex) until 9 weeks of age before behavioural experiments commenced (see below). One week after behavioural testing was completed, rats were killed by decapitation and their brains were collected and frozen on dry ice, then stored at −80°C until sectioning and processing for in situ hybridisation (see below).

### Preparation of MitoQ-loaded nanoparticles

Antioxidant MitoQ^42^ (a kind gift from Prof Mike Murphy, MRC Mitochondrial Biology Unit, Cambridge) was loaded onto γ-PGA-Phe nanoparticles (NP; a kind gift from Prof Mitsuru Akashi, Osaka University)^73^ to produce MitoQ-NP were made as previously described^31^. Soluble MitoQ or MitoQ-NP were used at a final dose of 0.5 μM MitoQ.

### Quantification of oxidative stress in placental, foetal and maternal tissues

Levels of reactive oxygen species (ROS) were measured in foetal, maternal and placental tissues (collected in Experiment 1) using the 2′,7′-dichlorofluorescein diacetate (DCFDA) (Sigma-Aldrich) assay. Sagittal sections (10 μm) were cut on the cryostat and exposed to 20 μM DCFDA solution in HBSS at 37 °C in a humidifying chamber. After counter-staining with DAPI, slides were immediately imaged using a confocal microscope (excitation: 495 nm, emission: 529 nm). Using Image-Pro Premier 9.2 (Media Cybernetics, USA), fluorescence levels of DCF, the product of DCFDA deacetylation by cellular esterases and oxidisation by ROS, were quantified.

### Preparation of placenta conditioned media

Conditioned tissue culture media was prepared using fresh rat placentae isolated from rats in Experiment 1 on GD20. Whole placentae (n = 3/litter) were incubated individually at 21% O_2_, 5% CO_2_ and 37°C in neuronal culture medium (Gibco Neurobasal medium supplemented with 1x Gibco B-27 supplement, 2 mM L-glutamine and 250 μM penicillin-streptomycin; all Thermo Fisher Scientific). After 24 h, the media around the explants was collected, syringe filtered (0.2 μm), frozen on dry ice and stored at −80°C until further use.

### Determination of plasma corticosterone concentrations

Plasma corticosterone concentrations in maternal plasma and foetal pooled plasma (collected in Experiment 1) were determined in duplicate using a commercially available radioimmunoassay kit (MP Biomedicals, Germany, sensitivity: 7.7 ng/ml, intra-assay variation 4.4–10.3%).

### Analysis of microRNA in placental conditioned medium and foetal plasma

The microRNA profile of the placental conditioned medium and foetal plasma samples (collected in Experiment 1) was analysed as described previously^31^. Briefly, total RNA was extracted from 200 μl conditioned medium or 100 μl foetal plasma using the miRNeasy Mini Kit and the miRNeasy Serum/Plasma Kit (Qiagen, Germany). microRNA expression levels were analysed using the nCounter Rat v1.5 miRNA Expression Assay (NanoString Technologies, USA), which detects over 400 different species-specific microRNAs. Undiluted samples (3 μl) were hybridised with barcoded probes and immobilised on an nCounter Cartridge. Barcode signals were counted using the nCounter Digital Analyser.

### Cortical cultures

Cortical cultures were prepared from dissociated rat embryonic day 18 cortical tissue and grown on glass coverslips as described previously^40^, in neuronal culture medium (see above). Placental conditioned medium or foetal plasma (n = 3, each replicate originating from a different dam; collected in Experiment 1) was applied to the cortical cultures for 6 days starting from day 12 *in vitro*. Exposures were performed at least in triplicate with 3 different sets of cortical cultures.

### Immunocytochemistry of cortical cultures

Cortical cultures were fixed in supercold methanol (−20 °C) and blocked with 5% bovine serum albumin, 5% normal goat serum in PBS for 30 min. Cultures were then incubated with primary antibodies against MAP2 (1:500, ab32454; abcam), tyrosine hydroxylase (1:500, ab112; abcam), GABA Aα1 (1:500, ab33299; abcam), GABA Aα2 (1:500, ab193311; abcam), GABA B1 (1:500, ab55051; abcam) or GluN1 (1:500, ab9864; Merck Millipore), overnight at 4°C. Sections were probed with secondary antibodies Alexa Fluor 488 anti-rabbit IgG or Alexa Fluor 488 anti-mouse IgG or (all Thermo Fisher Scientific, diluted 1:500) for 2 h at room temperature under minimal light conditions, washed with PBS and mounted in DAPI mounting media. Five images per coverslip were taken on a confocal microscope (SP2-AOBS, Leica). Dendrite lengths were measured using ImageJ. For the analysis of receptors, fluorescent images were taken at 64x magnification (with oil) using a Leica SP5II fluorescence microscope after excitation at 488 nm. Using ImageJ, images were converted to RGB files and measurements were taken of the mean grey value of each image providing an average of the relative intensity of the staining (as described previously^31^).

### Immunohistochemistry of P30 offspring brains

P30 brains collected in Experiment 2 were processed for immunohistochemistry. Three brains per group (one from each litter) were selected. Cryostat sections (12 μm) were cut and mounted as contiguous triplicates. The sections were fixed in cold methanol (−20 °C). For GluN1 staining, sections were fixed in 2% PFA, followed by permeabilisation in 0.3% Triton X-100 in PBS for 15 min. Sections were blocked with 5% goat serum, 0.3% Triton X-100 in PBS for 2 h at 4°C, followed by an overnight incubation (or 48 h incubation for GluN1 staining) at 4°C in primary antibody in PBS with 1% BSA, 0.3% Triton X-100. Primary antibodies were used against MAP2 (1:500, ab32454; abcam), tyrosine hydroxylase (1:500, ab112; abcam), parvalbumin (1:500, ab11427; abcam) GABA Aα1 (1:500, ab33299; abcam), GABA Aα2 (1:200, ab193311; abcam), GABA B1 (1:500, ab55051; abcam) or GluN1 (1:500, ab9864; Merck Millipore). Sections were incubated with secondary antibody Alexa Fluor 555 anti-rabbit IgG, Alexa Fluor 488 anti-mouse IgG or Alexa Fluor 568 anti-mouse IgG (Thermo Fisher Scientific) at 1:500 for 2 h at 4°C. Vectashield Mounting Medium with DAPI (Vector Laboratories, USA) was used to mount coverslips. Coronal sections were chosen to show the thalamic reticular nucleus, hippocampal regions (CA1, CA2 and CA3), the basolateral amygdala and, as control areas, cortical regions (primary somatosensory, primary auditory and retrosplenial granular cortex). For each region, in each brain, a minimum of 5 fields of view were examined for each of 3 sections in both hemispheres using a LASX (Leica) widefield microscope or a SP5II (Leica) confocal microscope at 40x magnification using oil. This resulted in a minimum of 30 fields of view being analysed for each brain region. The analysis of the intensity of the immunostaining of GluN1 and GABA receptor subunits was performed using a similar approach to that previously described^31^. Briefly, nine sections per treatment were imaged and 10-12 images were collected for every brain region using the same microscopy settings of intensity and magnification to allow for a meaningful comparison. Images were converted to greyscale and the total pixel number was determined using a corrected macro for ImageJ. The total pixel number was subsequently subtracted from the background.

### Western blot of P30 offspring brains

Total protein was extracted from foetal (Experiment 1) and P30 frontal cortex tissue (collected in Experiment 2) using RIPA buffer containing cOmplete Mini Protease Inhibitor Cocktail tablets (Sigma-Aldrich). Protein concentrations were quantified using the Pierce BCA Protein Assay Kit (Thermo Fisher Scientific). 20 µg of each sample was separated on a 10% SDS-PAGE gel and transferred onto a nitrocellulose membrane. Membranes were blocked in 5% milk in TBS-T and incubated overnight at 4°C with the primary antibodies anti-tyrosine hydroxylase (1:1,000; AB152; Merck Millipore) or anti-dopa decarboxylase (1:500; AB1569; Merck Millipore), followed by 1-h incubation with secondary antibody goat anti-rabbit Alexa Fluor 680 (1:10,000; A21108) or Alexa Fluor 800 (1:10,000; A32730; Thermo Fisher Scientific). Blots were stripped with Restore Plus Stripping Buffer (Thermo Fisher Scientific) prior to 1-h incubation with loading control antibodies anti-alpha-tubulin (1:1,000; ab15246; abcam) or anti-GAPDH (1:1,000; ab8245; abcam). Protein bands were visualised on the Odyssey Imaging System (LI-COR Biotechnology, Cambridge, UK).

### In situ hybridisation (ISH)

Adult offspring brains collected after behavioural testing in Experiment 2 were sectioned through the amygdaloid complex at 15 µm using a cryostat and thaw-mounted onto Polysine slides. Sections were then fixed, dehydrated and processed for ISH as previously described^4,61^. ^35^S-UTP radio-labelled (Perkin Elmer UK Ltd., Beaconsfield, Buckinghamshire, UK) cRNA sense and antisense probes were synthesised from the linearised pBluescript vector expressing a 518 bp cDNA fragment encoding rat *Crh* and used to detect *Crh* mRNA expression, as previously described^4,61^. Following overnight hybridisation and subsequent washes, slides were dipped in autoradiographic emulsion (Illford K5, Knutsford, Cheshire, UK), exposed at 4°C for 4 weeks, before being developed, counterstained with haematoxylin and eosin, and coverslipped. Sections hybridised with sense probes served as negative controls and showed no signal above background. For quantitative analysis, the region of interest (central amygdala) was localised under a light microscope and the number of cells expressing the mRNA of interest were manually counted bilaterally. A positive cell was defined as one with <5x the number of overlying silver grains than background.

### RNA sequencing

The transcriptome of the foetal frontal cortex (n = 4/treatment, originating from different litters; collected in Experiment 1) was analysed as previously described^31^. Briefly, RNA was extracted from 50 mg of foetal frontal cortex using the RNeasy Mini kit (Qiagen) and subjected on-column incubation with RNase-free DNase according to manufacturer’s protocol (Qiagen). RNA quality and integrity was measured on the 2100 Bioanalyzer (Agilent Technologies). mRNA sequencing was performed by Edinburgh Genomics, UK. Libraries were prepared from total RNA samples using the Illumina TruSeq stranded mRNA Sample Preparation Kit and sequenced by 75 bases paired-end sequencing across two lanes of an Illumina HiSeq 4000. This level of sequencing produced greater than 42 million paired reads per sample. The FASTQ files were generated using the standard Illumina pipeline for bcl2fastq.

### Bioinformatic analyses

Tophat^74^ was used to align RNA sequencing reads to the rat reference genome Rnor_6.0 (GenBank Assembly ID GCA_000001895.4). Total alignment rates were 83-85% on average, after removing multiple alignments and discordant pairs. Read counts were generated from the resulting BAM files using HTSeq^75^. Differential expression analysis was performed using DESeq. Genes were classed as significant differentially expressed genes for *p* < 0.05. Pathway analysis was performed using PANTHER 11.0^76,77^.

NanoString nCounter data was analysed using a pipeline described previously^31^. RUVSeq^78^ was used to adjust the counts to account for unwanted variation and then edgeR^79^ to predict differentially abundant microRNAs from the adjusted counts. miRNAs were classed as significant differentially secreted miRNAs if *p* < 0.05. Predicted targets of differentially abundant microRNAs were derived from TargetScanHuman v7.0^80^ (Total Context Score < −0.2)and subjected to gene ontology analysis using DAVID 6.8^81,82^. Enrichment analyses were performed in R/Bioconductor using the Fisher’s exact test and investigating enrichment only. Correlation of abundance changes of significant miRNAs with abundance changes of significant mRNAs was analysed with the miRComb package^83^ for R/Bioconductor, using Spearman correlation and Benjamini-Hochberg adjustment for multiple comparisons. Correlation analysis between log2 fold change values of prenatal stress exposure compared to control and prenatal stress exposure + MitoQ compared to controls was performed using Pearson’s correlation in R/Bioconductor. Venn diagrams were produced using Venny 2.1 (http://bioinfogp.cnb.csic.es/tools/venny/index.html).

### Behavioural tests on adult offspring

Starting between 9-10 weeks of age, offspring (one male and one female per litter; n = 8/group/sex; generated in Experiment 2) were subjected to a battery of behavioural tests. Anxiety tests were carried out first, starting with the light-dark box^84^ and followed by the elevated plus maze^85^ 4 days later. In the subsequent 2 weeks, the rats were tested for the presence of two hallmark traits of depression: anhedonia, using the sucrose preference test, and behavioural despair using the forced swim test^86^. A separate cohort of female offspring (aged 16-17 weeks; littermates of those used in the anxiety/depression tests) was tested in a social olfactory memory paradigm, as it was shown previously that prenatally stressed females, but not males, showed deficits in social odour memory^5^. In all tasks, the mazes were thoroughly cleaned with 70% ethanol and allowed to air-dry between each trial and males and females were tested on separate days.

#### Light-dark box

The light-dark box consisted of a transparent box and lidded opaque black box connected by an opening, as described previously^87^. Rats were placed into the dark chamber and allowed to explore freely for 5 min. The trials were recorded with an infrared camera and the time spent in each compartment and the latency to enter the light box was automatically scored using EthoVision XT software v12 (Noldus, Wagenigen, The Netherlands).

#### Elevated plus maze

The elevated plus maze consisted of 2 open arms, located opposite each other, intersected with 2 closed arms, elevated ca. 70 cm above the floor^87^. Rats were placed at the intersection, facing an open arm, and were allowed to explore freely for 10 min. The latency to enter the open arm, time spent in each arm, and the number of open arm entries was automatically quantified using EthoVision XT software.

#### Sucrose preference test

Rats were singly housed in clean IVCs with two drinking bottles per cage. Fluid consumption was measured by weighing the drinking bottles every 24 h over 3 days. On the first day, both bottles contained filtered tap water. On the second day, one water bottle was substituted for a bottle containing 2% (w/v) sucrose solution. The following day, the positions of the water and sucrose bottles were reversed. Preference for sucrose was expressed as a percentage, calculated by the volume of sucrose solution consumed, divided by the total volume of fluids consumed within the 48 h choice period.

#### Forced swim test

The forced swim test consisted of 2 swimming bouts spaced 24 h apart. On the first day, rats were forced to swim for 10 min in a 50 cm tall cylinder filled with water (23-25°C) to a depth of 30 cm. The following day, the rats were placed in the same cylinder, forced to swim for 5 min and their behaviour was digitally recorded. Videos were analysed using a pharmacologically validated time-sampling technique where the 5-min bout was divided into 60 x 5 s bins, and the predominant behaviour in each bin was recorded. Behaviours analysed included climbing (with vertical movement of forepaws against the walls of the cylinder), swimming (horizontal movement in the cylinder) and floating (assuming a passive position without struggling, making only small movements necessary to keep their heads above water).

#### Social olfactory memory

Wooden beads (25 mm in diameter) were impregnated with social odour by being kept for 1 week in a cage housing unrelated and unfamiliar female conspecifics. Rats were allowed to habituate to the test arena, a clean open top cage without bedding, for 2 h. In the exposure session, rats were first introduced to a wooden bead (designated as ‘familiar’) and were allowed to freely interact with it for 4 min. After a 3-h consolidation period, spent in the test arena cleaned of odours, the rats were exposed to 2 beads simultaneously for 4 min, a fresh familiar bead with the same odour as before and a bead with a novel odour. The time spent interacting with each bead (which included sniffing, biting or holding bead in mouth) was scored manually using Observer XT software (Noldus). The preference score was calculated as time spent in investigating the novel bead divided by the total time spent investigating both beads during the test session.

### Statistics

Analysis of behaviour and microscopy images was performed with the experimenter blind to the group allocation. Data are presented as group means ± s.e.m. As indicated in the figure legends, one-way or two-way ANOVA was performed in Prism 6.0 (GraphPad, USA) or SPSS 21.0 (IBM Corp., USA) with post-hoc analysis using Bonferroni correction, Tukey’s test or Student Newman Keuls test for multiple comparisons. Two-way ANOVA was used to test for main effects of drug and of stress conditions and for interaction effects.

### Data availability

Raw and processed data files generated from NanoString analysis and RNA sequencing have been deposited in NCBI’s Gene Expression Omnibus^88^ and are accessible through GEO Series accession number GSE131040.

## Acknowledgements

This work was supported by the BBSRC [grant number BB/J004332/1], The Perivoli Trust and The Waterloo Foundation [grant numbers I068-2946 and I890-3369]. Computing equipment was supported by the Engineering and Physical Sciences Research Council (EPSRC) [grant number EP/K008250/1]. The authors thank Prof Tudor Fulga at the University of Oxford for access to his NanoString nCounter system and Dr Bruno Steinkraus for technical help with the NanoString experiments.

## Author contributions

CPC and PJB contributed to study conception and design. HS, TJP, SY, AA and PJB performed the experiments. HS, TJP, SY, MFR, PJB and CPC analysed and interpreted data. HS, CPC and PJB wrote the manuscript. All authors critically read and approved the final version of the manuscript.

## Competing interests

HS and TJP have previously consulted for Placentum Ltd. The University of Bristol has filed a patent application for the nanoparticle formulation used in this study and its application to pregnancy-related diseases. The other authors report no conflict of interest.

