## Supplementary Information for "Maternal antioxidant treatment prevents behavioural and neural changes in offspring exposed to prenatal social stress"

#### Supplementary Data

**Supplementary Data S1 | Differentially expressed genes in the foetal cortex following exposure to prenatal social stress and maternal antioxidant treatment.** Cortical tissue from foetuses that had been exposed to prenatal social stress and maternal injection of MitoQ-NP was analysed using RNA sequencing. Data is presented in a Supplementary Excel file. Differentially expressed genes ($p < 0.05$, fold change ≥ ±0.25) between non-stressed controls with vehicle injection and prenatally stressed rats with vehicle injection are shown on Sheet 1. Sheet 2 shows the differentially expressed genes between prenatally stressed rats treated with vehicle and those treated with MitoQ-NP.

**Supplementary Data S2 | Differentially abundant microRNAs in the foetal plasma and placenta-conditioned medium following exposure to prenatal social stress and maternal antioxidant treatment.** Foetal plasma and culture medium conditioned by placentae collected from pregnant rats that had been exposed to prenatal social stress with and without maternal administration of MitoQ-NP was analysed for microRNA expression levels. Data is presented in a Supplementary Excel file. Differentially expressed genes ($p < 0.05$) between non-stressed administered vehicle and prenatally stressed rats administered vehicle are shown on Sheet 1 for foetal plasma and in Sheet 2 for placenta-conditioned medium. Sheet 3 and 4 show the differentially expressed genes between prenatally stressed rats with vehicle injection and those with MitoQ-NP injection for foetal plasma and placenta-conditioned medium, respectively.

**Supplementary Figure legends**

**Supplementary Figure S1 | Behavioural effects of prenatal social stress and maternal antioxidant treatment on juvenile offspring.** Male and female juvenile offspring of rats exposed to social stress and maternal injection of vehicle or MitoQ-NP were tested in the light/dark box (a,e), the elevated plus maze (b,f) and the sucrose preference test (c,g). Two-way ANOVA revealed a significant effect of MitoQ-NP on the total distance travelled in the light/dark box in males ($F_{1,27}$ = 5.2, $p$ = 0.030; a) and a significant stress x MitoQ-NP interaction in females ($F_{1,27}$ = 4.34, $p$ = 0.047; e). On the elevated plus maze, there was a significant effect of stress on the number of entries into open arms in male rats ($F_{1,27}$ = 4.82, $p$ = 0.037; b) and a significant stress x drug interaction on total entries into all arms in female rats ($F_{1,27}$ = 9.24, $p$ = 0.005; f). There was no effect of prenatal stress of MitoQ-NP treatment on sucrose consumption in either males (c) or females (g). *Crh* mRNA expression was measured in the central amygdala of male rats on postnatal day 30 (P30) and showed a significant stress x drug interaction ($F_{1,27}$ = 3.41, $p$ = 0.080; d). Female offspring were further tested in a social olfactory memory task and the time spent exploring the novel odour or the familiar odour is shown (h). *$p < 0.05$; **$p < 0.01$.

**Supplementary Figure S2 | Neuroanatomical effects of prenatal social stress and maternal antioxidant treatment in the brains of juvenile offspring.** Dendrite lengths (a,e), TH+ cell process lengths (b,f), TH+ cell counts (c,g) and PV+ cell counts (d,h) were quantified in juvenile male and female offspring born to mothers exposed to social stress and vehicle or MitoQ-NP injection during pregnancy. Auditory (ACTX), somatosensory (SCTX) and retrosplenial cortex (RCTX), along with thalamic reticular nucleus (TRN) were assessed. *$p < 0.05$, **$p < 0.01$, ***$p < 0.001$. $n$ = 6 (male), $n$ = 6 (female).

**Supplementary Figure S3 | Neuronal densities in the brains of juvenile offspring exposed to prenatal social stress and maternal antioxidant treatment.** Neuronal counts were assessed in male and female juvenile offspring exposed to prenatal social stress and maternal intravenous injection of vehicle or MitoQ-NP. CA1, CA2 and CA3 regions of the hippocampus (a,e), auditory (ACTX), somatosensory (SCTX) and retrosplenial cortex (RCTX) (b,f), basolateral amygdala (BLA) (c,g) and thalamic reticular nucleus (TRN) (d,h) were analysed. *$p < 0.05$, **$p < 0.01$, ***$p < 0.001$. $n = 6$ (male), $n = 6$ (female).

**Supplementary Figure S4 | Neurochemical effects of prenatal social stress and maternal antioxidant treatment in the brains of juvenile offspring.** The relative immunoreactivity for the GABA receptor subunits GABA Aα1 (a,e), GABA Aα2 (b,f) and GABA B1 (c,g) and for the glutamate receptor subunit GluN1 (d,h) were quantified in juvenile male and female offspring born to mothers exposed to social stress and vehicle or MitoQ-NP injection during pregnancy. Auditory (ACTX), somatosensory (SCTX) and retrosplenial cortex (RCTX), along with thalamic reticular nucleus (TRN) were assessed. *$p < 0.05$, **$p < 0.01$, ***$p < 0.001$. $n = 6$ (male), $n = 6$ (female).

### Supplementary Figures

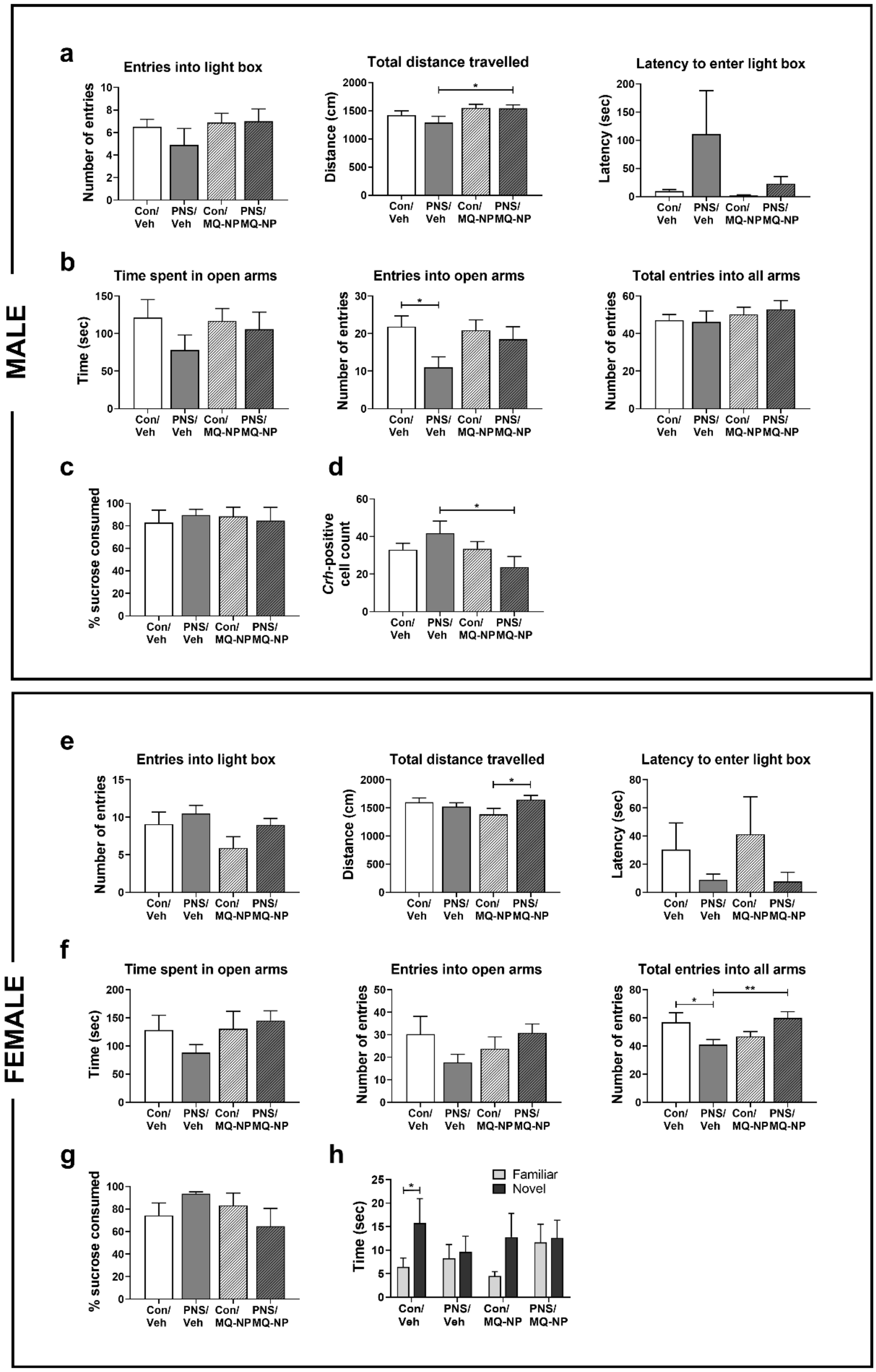


Supplementary Figure S1

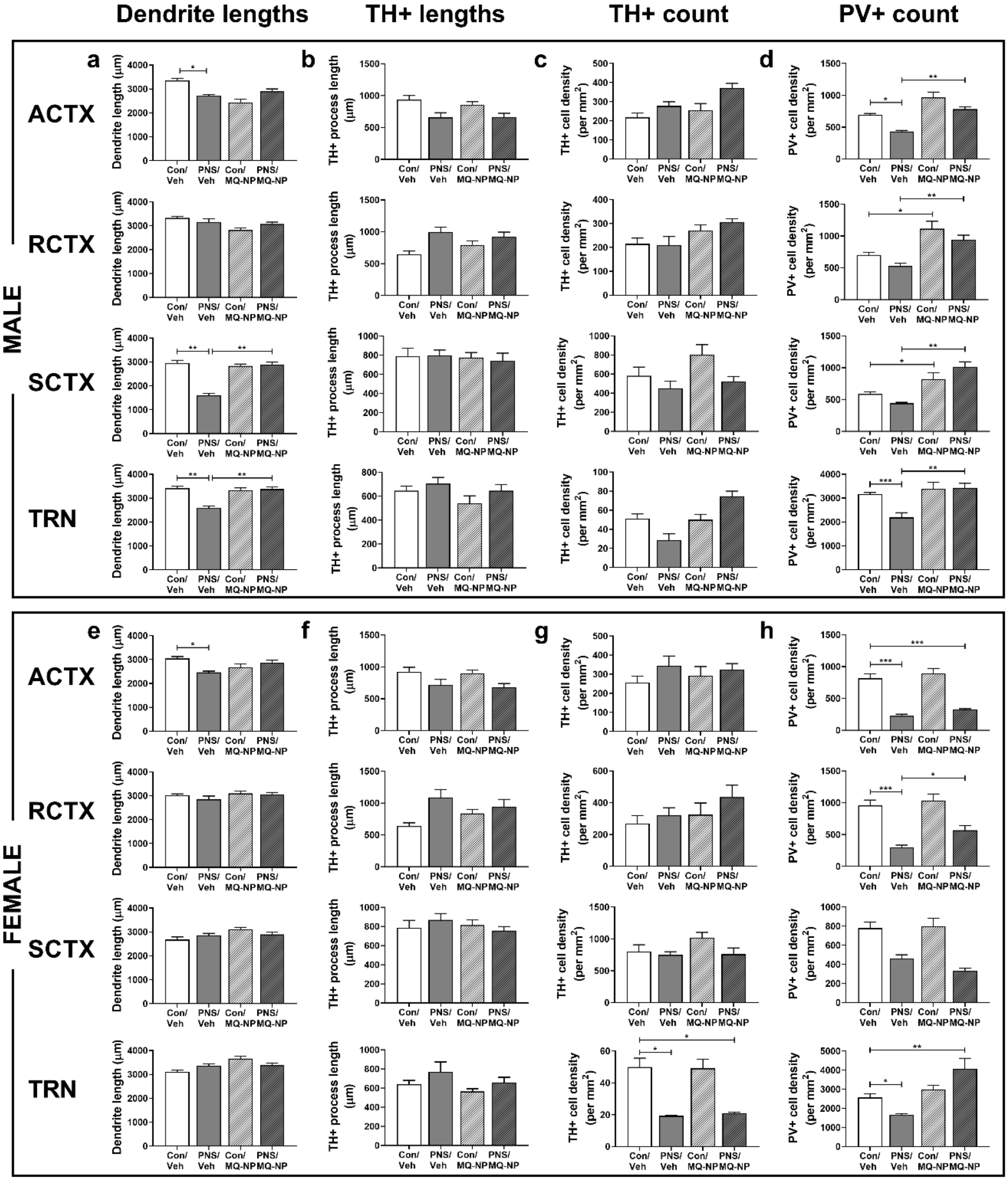


**Supplementary Figure S2**

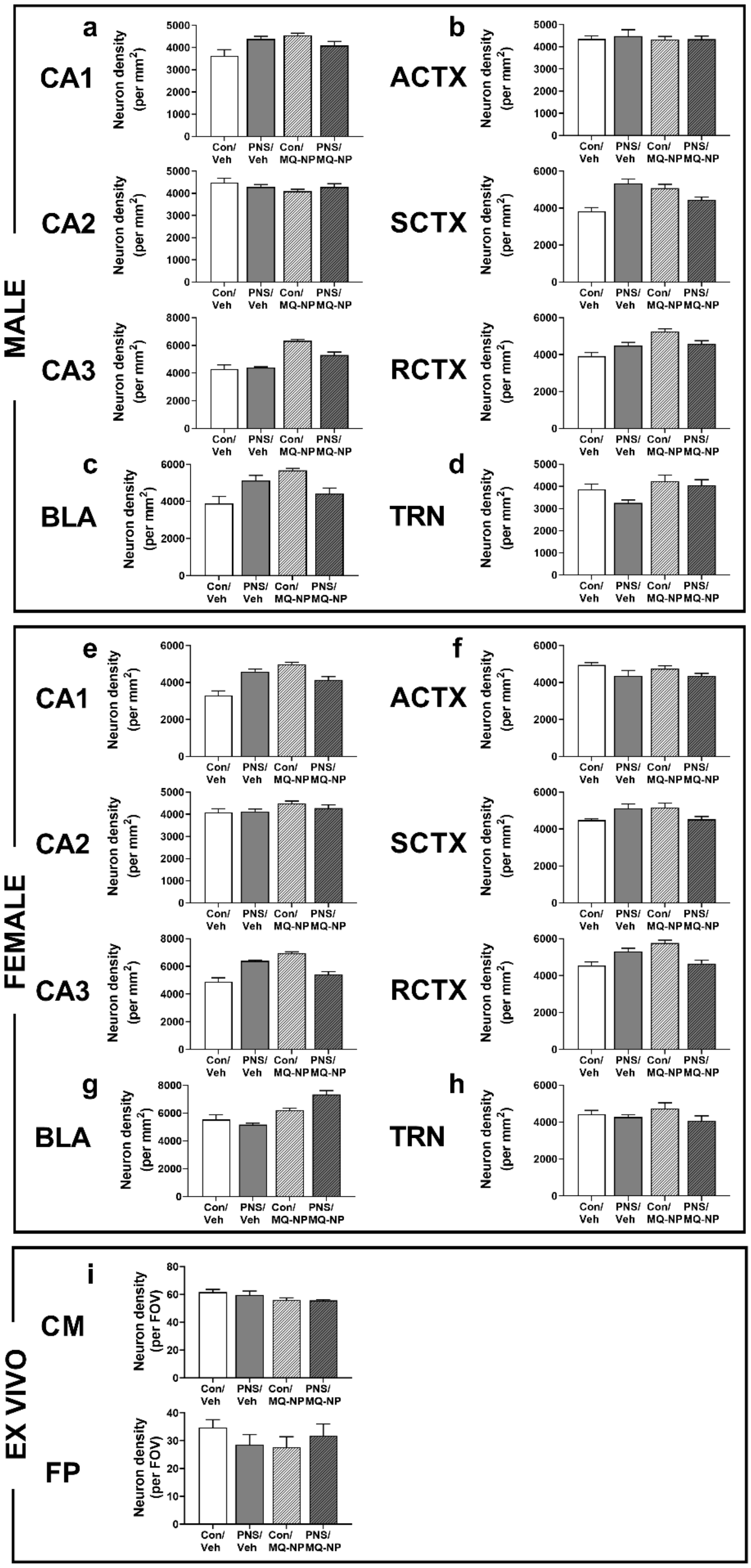


**Supplementary Figure S3**

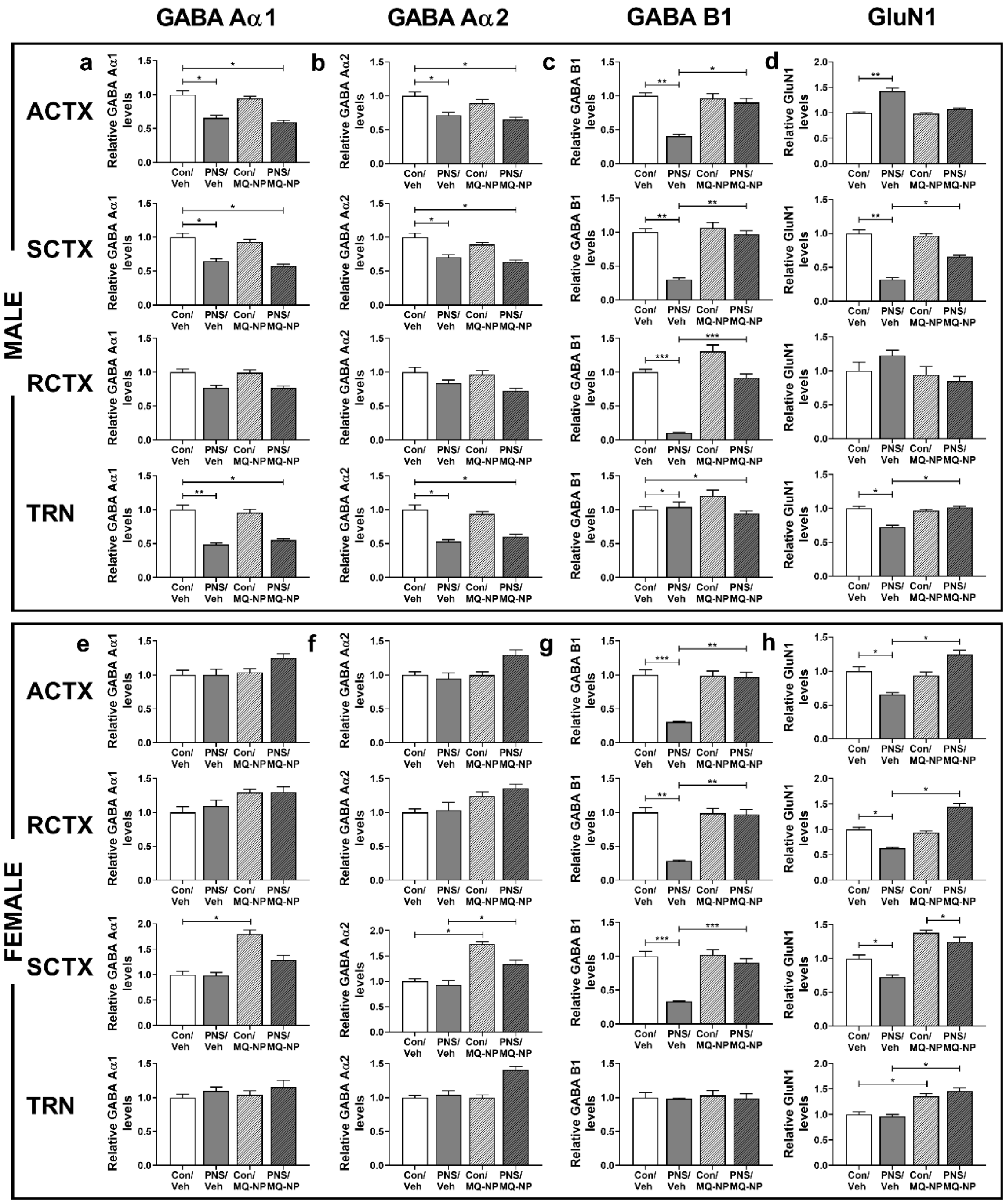


**Supplementary Figure S4**
